# Defective pollen meiosis in Arabidopsis due to combined arabinan and galactan insufficiency

**DOI:** 10.1101/2025.02.03.636199

**Authors:** Takuma Kikuchi, Kouichi Soga, Toshihisa Kotake, Daisuke Takahashi

## Abstract

The development of pollen is critical or seed plants, and depends on precise cellular and molecular mechanisms. It is known that the cell wall plays a key role in the progression of pollen maturation. An earlier phase in pollen development is the meiosis of pollen mother cells (PMCs), the fundamental process for producing viable pollen grains. However, the significance of the cell wall during pre-meiosis processes has remained unclear. Pectin, a major component of the cell wall, is known to accumulate abundantly in pollen. To investigate the significance of the cell wall during pollen meiosis, we generated an *arad1 gals2 gals3* triple mutant of *Arabidopsis thaliana*, lacking the genes responsible for normal synthesis of arabinan and galactan, which constitute the side chains of pectin. Although vegetative growth and cell wall properties were comparable in wild-type (WT) and *arad1 gals2 gals3*, the pollen development in the mutant failed during meiosis. Immunohistochemical analysis showed that pectic arabinan and galactan accumulated in WT PMCs before meiosis, but this was not observed in mutant PMCs. We believe that this study is the first to demonstrate the critical role of cell wall components, specifically pectic arabinan and galactan, in the pre-meiosis processes of pollen development.

**Highlight:** Pectic arabinan and galactan accumulate on the surface of pollen mother cells and this accumulation is essential for pollen meiosis.

## Introduction

The development of pollen is a pivotal event for reproduction in seed plants. Pollen development occurs in the anthers and is divided into 14 stages (Sanders *et al*., 1999; Scott *et al*., 2004; Alvarez-Buylla *et al*., 2010). Stages 1 to 5 of anther and pollen development take place before meiosis, while stages 6 to 14 involve pollen maturation during and after meiosis. Several cell wall components have been reported to play an important role during the maturation phases of pollen for successful pollen production (Worrall *et al*., 1992; Preuss *et al*., 1994; Enns *et al*., 2005; Blackmore *et al*., 2007; Ariizumi and Toriyama, 2011; Wan *et al*., 2011; Mollet *et al*., 2013; Cankar *et al*., 2014; Shi *et al*., 2015; Yin *et al*., 2022; Hasegawa *et al*., 2023) (Supplementary Table S2). Abnormalities in the synthesis of these components resulting in modification of cell walls during pollen maturation often lead to male sterility (Worrall *et al*., 1992; Chen *et al*., 2007; Begcy *et al*., 2019; Hasegawa *et al*., 2023).

The cell wall provides cells with strength, determines their shape and properties, and influences their ultimate development. Cell wall composition varies across plant organs, imparting distinct characteristics to each. In dicotyledons, the primary cell wall is mainly composed of three polysaccharides: pectin, hemicellulose and cellulose. Pectin and hemicellulose, known as matrix polysaccharides, serve as reinforcers that fill in and connect the cellulose microfibril network (Carpita and Gibeaut, 1993), and contribute to the mechanical properties and porosity of the cell wall (Fleischer *et al*., 1999; Peaucelle *et al*., 2011; Klaassen and Trindade, 2020). Pectin consists of three main domains: homogalacturonan (HG), rhamnogalacturonan-□ (RG-I), and RG-□. HG is a region of linear d-galacturonic acid (GalA) polymers that fill the gaps in the cell wall through calcium cross-linking, which depends on the degree of HG methylation (Tibbits *et al*., 1998). RG-II forms pectin network by cross-linking with boron (Matoh, 1997; Fleischer *et al*., 1999). RG-I is composed of a main chain having repeating sequences of GalA and d-rhamnose (Rha), with side chains of linear β-1,4-galactan composed of d-galactose (Gal), branched α-1,5-arabinan composed of l-arabinose (Ara), and type-□ arabinogalactan composed of both Gal and Ara attached to the Rha residue (Kaczmarska *et al*., 2022). Pectic arabinan, in particular, is involved in mechanisms such as stomatal opening and closing, cell wall flexibility in inflorescence stems, and pollen tube elongation (Jones *et al*., 2003; Verhertbruggen *et al*., 2013; Lampugnani *et al*., 2016; Carroll *et al*., 2022). On the other hand, pectic galactan is associated with environmental stress responses such as the adaptation process to salt and freezing stress that cause its accumulation (Wang *et al*., 2021; Takahashi *et al*., 2024).

During pollen development and maturation, cell wall composition and structure differ from what is seen during vegetative growth. For example, mature pollen cell walls consist of two layers: the inner (intine) and the outer (exine). The intine contains layers of pectin, cellulose, and callose (Ma *et al*., 2021). In particular, callose, a linear β-1,3-glucan, is a major component of the pollen cell wall during maturation, and its eventual degradation and utilization is an essential factor in the final maturation of pollen (Dong *et al*., 2005; Chen and Kim, 2009). If callose degradation is disrupted, pollen grains develop abnormal morphology, as seen in *Oryza sativa* and *Nicotiana tabacum* (Worrall *et al*., 1992; Chen and Kim, 2009; Wan *et al*., 2011). This indicates that cell wall components like callose play essential physiological roles in the post-meiotic pollen maturation process.

Pectin, along with callose, has also been recognized as a crucial component of the cell wall involved in pollen development and maturation (Kandasamy *et al*., 1994; Bárány *et al*., 2010). Recent studies have highlighted that the degree of methylesterification of HG is essential for both pollen development and maturation (Hasegawa *et al*., 2023). Pectic arabinan side chains are localized in pollen cell walls and play a significant role in pollen maturation, especially in potato (Dardelle *et al*., 2010; Castro *et al*., 2013; Cankar *et al*., 2014). Similarly, galactan accumulates in the cell wall of mature pollen in olive and potato, although it is thought to be less involved in pollen maturation (Castro *et al*., 2013; Cankar *et al*., 2014). Pectic arabinan and galactan are present as neutral sugar side chains in the RG-I region of pectin and may therefore have similar functions in certain physiological processes. However, their specific roles in pollen development and maturation remain largely unknown.

We thus decided to investigate the roles of pectic arabinan and galactan in pollen development in the model plant Arabidopsis (*Arabidopsis thaliana*). Arabinan is synthesized by two enzymes: ARABINAN DEFICIENT 1 (ARAD1) and ARAD2 (Harholt *et al*., 2006, 2012), while galactan synthesis in Arabidopsis is carried out by three enzymes: GALACTAN SYNTHASE 1 (GALS1), GALS2, and GALS3 (Liwanag *et al*., 2013; Ebert *et al*., 2018). Despite the low level of arabinan in *arad1* and *arad2* mutants, and the low level of galactan in the *gals1 gals2 gals3* triple mutant, all these mutants still produce seeds comparable to wild type (WT). Therefore, the roles of pectin, and particularly arabinan and galactan, in Arabidopsis pollen development have so far remained unclear.

To further examine these roles, we generated an *arad1 gals2 gals3* triple mutant with reduced levels of both arabinan and galactan in pectin side chains and observed its pollen development. Our results demonstrated that simultaneous reduction of pectic arabinan and galactan causes complete male sterility due to abnormal pollen development. Detailed observations of the pollen development revealed that the polysaccharides in question accumulate on the pollen mother cells (PMCs) surface just before meiosis, and that abnormal morphology of PMCs manifests during meiosis in the mutants. Although several studies have previously shown that cell wall polysaccharides are involved in pollen maturation, this study focuses on their importance in the early stages of pollen development before meiosis.

## Materials and Methods

### Plant materials and growth conditions

WT Arabidopsis Columbia-0 seedlings were sown on Murashige and Skoog agar medium containing 2% sucrose and grown for one week at 22°C under a 16-hour light period (120 µmol m^-2^ s^-1^) followed by an 8-hour dark period. Subsequently, they were transplanted into rock wool and grown under the same conditions. The plants used were *arad1* mutant (SALK_029831) (Harholt *et al*., 2006), lacking *ARAD1*, related to arabinan synthesis, and *gals1 gals2 gals3* triple mutant (SALK_016687, SALK_121802, WiscDsLox377-380G, respectively) (Ebert *et al*., 2018), lacking *GALS1*, *GALS2* and *GALS3*, related to galactan synthesis. Stamens and pistils of the *arad1* mutant and *gals1 gals2 gals3* triple mutants were used for crossing. The resulting seeds were grown and screened by PCR using designated primers (Supplementary Table S1) to confirm the *arad1 gals2 gals3* triple mutation.

### Extraction of the pectic polysaccharides and sugar composition analysis

The plant parts except for the roots were ground using a mortar and pestle, then centrifuged at 21,500 × *g* at 4°C for 5 minutes. The supernatant was collected to remove soluble sugars. The precipitates were suspended in 800 µL of 80% ethanol, heated at 100°C for 2 minutes, and centrifuged at 21,500 × *g* at 4°C for 5 minutes. The supernatant was collected to remove lipids and pigments. Then, 500 µL of water and 500 µL of amylase reaction solution (50 mM MOPS-KOH, pH 6.5; amylase, 20 units/mL, SIGMA) were added to the precipitate, shaken at 37°C for 2 hours, centrifuged at 21,500 × *g* at 4°C for 5 minutes, and the supernatant was discarded to remove starch. After centrifugation at 21,500 × *g* at 4°C for 5 minutes, the supernatant was collected as the hot water fraction (hot water fr.) rich in low molecular weight pectin. These steps were repeated twice. Then, 500 µL of EDTA solution (50 mM Na-Phosphate, pH 6.8; 50 mM EDTA-2Na, pH 8.0) was added to the precipitate and suspended, boiled at 100°C for 10 minutes, centrifuged at 21,500 × *g* at 4°C for 5 minutes, and the supernatant was collected. These steps were repeated twice to obtain an EDTA fraction (EDTA fr.) rich in high molecular weight pectin.

The EDTA fractions obtained by fractionation were dialyzed with water for 2 days and both hot water and EDTA fractions were lyophilized. The extracted cell wall polysaccharides (50 µg) were dissolved in 100 µL of water and 100 µL of 4 N trifluoroacetic acid (TFA) was added to a final concentration of TFA of 2 N. Subsequently, they were hydrolyzed by heating at 120°C for 60 minutes. TFA was removed by drying in a Speed Vac (TAITEC VC-36R, Japan) and the sample was dissolved in 250 µL of water after triple centrifugal concentration with 50 µL of water. The 50 µL portion was used for sugar composition analysis by high-performance anion exchange chromatography-pulsed amperometric detection (HPAEC-PAD) with the ICS-5000+ series system equipped with a CarboPac PA-1 column (Thermo Fisher Scientific, USA). Elution was carried out with water, 0.1 M sodium hydroxide and 0.1 M sodium hydroxide with 0.5 M sodium acetate at a flow rate of 1.0 mL/min at room temperature (RT) as described previously (Ishikawa *et al*., 2000).

### Morphological observation of bud interior

Non-flowering buds of WT and *arad1 gals2 gals3* triple mutants grown for more than 50 days were collected. Fixing solution (1.75 M acetic acid in ethanol) was added, vacuumed in a desiccator, and allowed to stand at RT for 2 hours. The fixing solution was removed and subsequently 90% ethanol was added, vacuumed, and allowed to stand at RT for 20 minutes. These steps were repeated with 70% ethanol, 50% ethanol, 30% ethanol and water. Transparency solution (40 g chloral hydrate in 5 mL glycerol and 10 mL water) was added and allowed to stand overnight at RT. Then, the transparent buds were observed by a stereomicroscope (S9i, Leica Microsystems, Germany).

### Observation of pollen by immunohistochemical staining

Plants were grown for more than 50 days, and non-flowering buds were collected. Fixing solution (10 mL 4% paraformaldehyde; 250 µL 25% glutaraldehyde) was added, the buds were vacuumed in a desiccator and allowed to stand at RT for 6 hours. The fixing solution was removed and 1×PBS was added to the samples. Then, samples were vacuumed and allowed to stand at RT. The same steps were repeated with the following solutions: 10% ethanol (6 hours), 30% ethanol (12 hours), 50% ethanol (12 hours), 70% ethanol (12 hours), 80% ethanol (12 hours), 90% ethanol (12 hours), 100% ethanol (12 hours) and 100% ethanol (12 hours). A mixture of Technovit solution (100 mL Technovit 7100 supplemented with 1 g Hardener □) (Kulzer Technic, Germany) and ethanol (Technovit: ethanol = 1: 5) was added and samples were allowed to stand for 6 hours at 4°C. The same steps were repeated with the following solutions: Technovit: ethanol = 1:3 (12 hours), 3:2 (12 hours), 5:1 (12 hours), 1:0 (12 hours) and 1:0 (12 hours). After Technovit replacement, embedding solution (1 mL Technovit solution supplemented with 100 µL Hardener □) were added and allowed to stand for 12 hours at 4°C. The samples were then incubated at 60°C for 1 hour for curing.

Embedded samples were cut and sectioned using a rotary microtome (Leica, MR2125RT, Germany). Sections were adhered to MAS glass slides (MATSUNAMI, MAS-01, Japan) and used as samples for microscopy. Blocking buffer containing 3% (w/v) skim milk in PBS (0.137 M NaCl; 2.7 mM KCl; 10 mM Na_2_HPO_4_; 1.8 mM KH_2_PO_4_, pH 7.4) was added to the sections on the glass slides and allowed to stand for 30 minutes. The blocking buffer was removed, and 3 µL each of LM5 and LM6 antibodies (diluted 5-fold in blocking buffer, Kerafast, USA), which recognize pectic arabinan and galactan, respectively, were added and allowed to incubate for 2 hours in the dark at RT. The primary antibody was then removed and washed three times with PBS. Subsequently, 3 µL of secondary antibody (Alexa Fluor 488, Thermo Fisher Scientific, USA) was added and allowed to stand for 1 hour in the dark at RT. The secondary antibody solution was then removed and washed three times with PBS. In addition, 3 µL of calcofluor white (2.5 mg calcofluor white in 10 mL PBS) was added and samples were allowed to stand for 10 minutes in the dark at RT. The samples were then washed three times with PBS. The sections were observed by an epifluorescence microscope (ECLIPSE Ci-L Plus, Nikon, Japan).

For toluidine blue and aniline blue staining, sections obtained above were stained with 0.05% toluidine blue (Waldeck GmbH, Germany) in water (1 minutes at RT) and 0.01% aniline blue (FUJIFILM Wako Pure Chemical, Japan) in 77 mL phosphate buffer (10 minutes at RT), respectively. The sections were then washed three times with water and observed by an epifluorescence microscope.

## Results

### The reduction of arabinan and galactan affects seed formation

We generated an *arad1/+ gals1/+ gals2/+ gals3/+* heterozygous mutant by crossing *arad1* mutant with *gals1 gals2 gals3* triple mutant. Subsequently, we selected for plants having homozygous T-DNA insertions in three genes using PCR from seeds obtained by self-pollination of heterozygous mutants, yielding seedlings of *arad1 GALS1 gals2 gals3* triple mutant (*arad1 gals2 gals3*, Fig. 1A). As the chromosomal positions of *ARAD1* and *GALS1*are very close to each other (14793567 - 14795701 bp and 14216771 - 14219252 bp regions on chromosome 2, respectively), *arad1* and *GALS1* are genetically linked and the obtained *arad1 gals2 gals3* did not have a mutated *GALS1* gene.

**Fig. 1.**
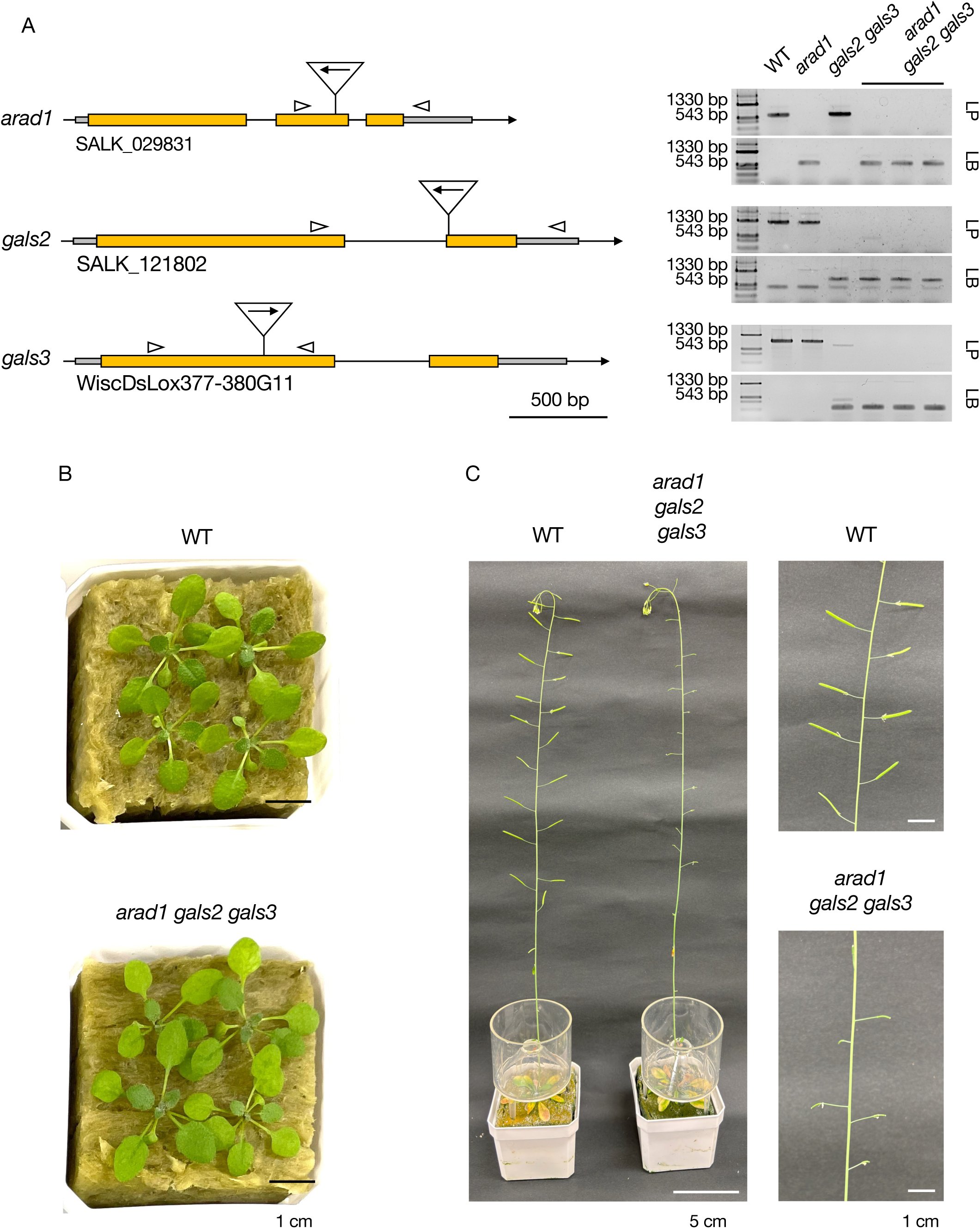
Verification of T-DNA insertion in *ARAD1*, *GALS2*, and *GALS3* genes and phenotypes of *arad1 gals2 gals3*. (A) The T-DNA insertion sites for *ARAD1*, *GALS2*, and *GALS3* are indicated with arrows, where the direction of the arrows represents the orientation of T-DNA insertion. The arrowheads mark the binding sites of the primers used in this study (Supplementary Table S1). Scale bar = 500 bp. The figure on the right shows the results of PCR, indicating that T-DNA insertion has occurred in the genes corresponding to each mutant. LP represents the PCR performed with the left primer (LP) and right primer (RP), while LB represents the PCR performed with the T-DNA left border primer (LB) and RP. (B) Phenotypic observation of rosette leaves in WT and *arad1 gals2 gals3* of seedlings for 3 weeks. Scale bar = 1 cm (C) Phenotypic observation of inflorescence stem and siliques in WT and *arad1 gals2 gals3* of seedlings after more than 7 weeks. Scale bar = 5 cm and 1 cm. Alt text: (A) A schematic structure of the gene sequences for *arad1*, *gals2* and *gals3*. Below the gene sequences are the accession numbers of the mutants used in this study. On the right, genotyping results are shown, indicating that T-DNA was inserted in *ARAD1*, *GALS2* and *GALS3*. We also separately confirmed that these mutants do not contain the WT allele. (B) The three-week-old seedlings of WT and *arad1 gals2 gals3* were observed, confirming that *arad1 gals2 gals3* does not exhibit morphological changes in the rosette leaves compared to WT. (C) Phenotypic observation of inflorescence stem and siliques in WT and *arad1 gals2 gals3*. The right side shows an enlarged image of the siliques, demonstrating that, in *arad1 gals2 gals3*, the siliques failed to develop, and no seeds were formed.

First, we investigated the composition of the cell wall polysaccharides in the leaves of both WT and *arad1 gals2 gals3*. The *arad1 gals2 gals3* leaves showed no significant decrease in total pectin content (Supplementary Fig. S1A). However, the levels of Ara and Gal were significantly reduced, with respective decreases of 30% and 25% in mol% composition, and 31% and 27% in sugar content compared to WT (Supplementary Figs. S1B, C). To see if these components affected the mechanical properties of the cell wall, extensibility and breaking force of the cell wall in leaves were also measured, but *arad1 gals2 gals3* as well as *arad1* and *gals2 gals3* were not significantly different from the WT (Supplementary Fig. S2). Indeed, the reduction of Ara and Gal in the pectin did not affect plant morphology at the vegetative growth stages (Fig. 1B). However, extremely small siliques were observed in *arad1 gals2 gals3* compared to WT (Fig. 1C).

To examine whether these phenotypes were linked to genotype, seeds produced by self-pollinating *arad1/+ gals1/+ gals2 gals3* heterozygous plants were grown. Subsequently, the genotype of each plant was determined by PCR and the number of seeds per silique was counted (Supplementary Fig. S3). The results showed that all *arad1 gals2 gals3* plants were seedless, while *arad1/+ gals2 gals3* had a slightly lower average seed count compared to *gals1 gals2 gals3*. This indicates that both pectic arabinan and galactan were involved in the gametogenesis, fertilization, and/or seed formation processes.

### Arabinan and galactan are involved in pollen formation

We therefore tried to determine the reason for the inability of arad1 gals2 gals3 to produce seed and investigated anthers and pistils under a stereomicroscope. In contrast to WT (Fig. 2A-C), anthers did not form or were very short and small in *arad1 gals2 gals3* even though bud development progressed (Figs. 2D-F). Therefore, to examine the formation of anthers in the early stages of floral development, the bud tissue was made transparent, and internal structures were observed with a stereomicroscope (Figs. 2G-L). Yellow parts derived from exine, a specialized cell wall structure formed on the pollen surface, was observed in WT anthers, whereas no such structures were observed in *arad1 gals2 gals3*. On the other hand, pistils were observed in *arad1 gals2 gals3* as well as in WT, indicating that the *arad1 gals2 gals3* mutation affects pollen development in the anthers, causing male sterility but does not impinge on pistil development. Indeed, when WT pollen was transferred to *arad1 gals2 gals3* pistils, the pistils successfully formed seeds and PCR confirmed that they had inherited the genes from the WT (Supplementary Fig. S4).

**Fig. 2.**
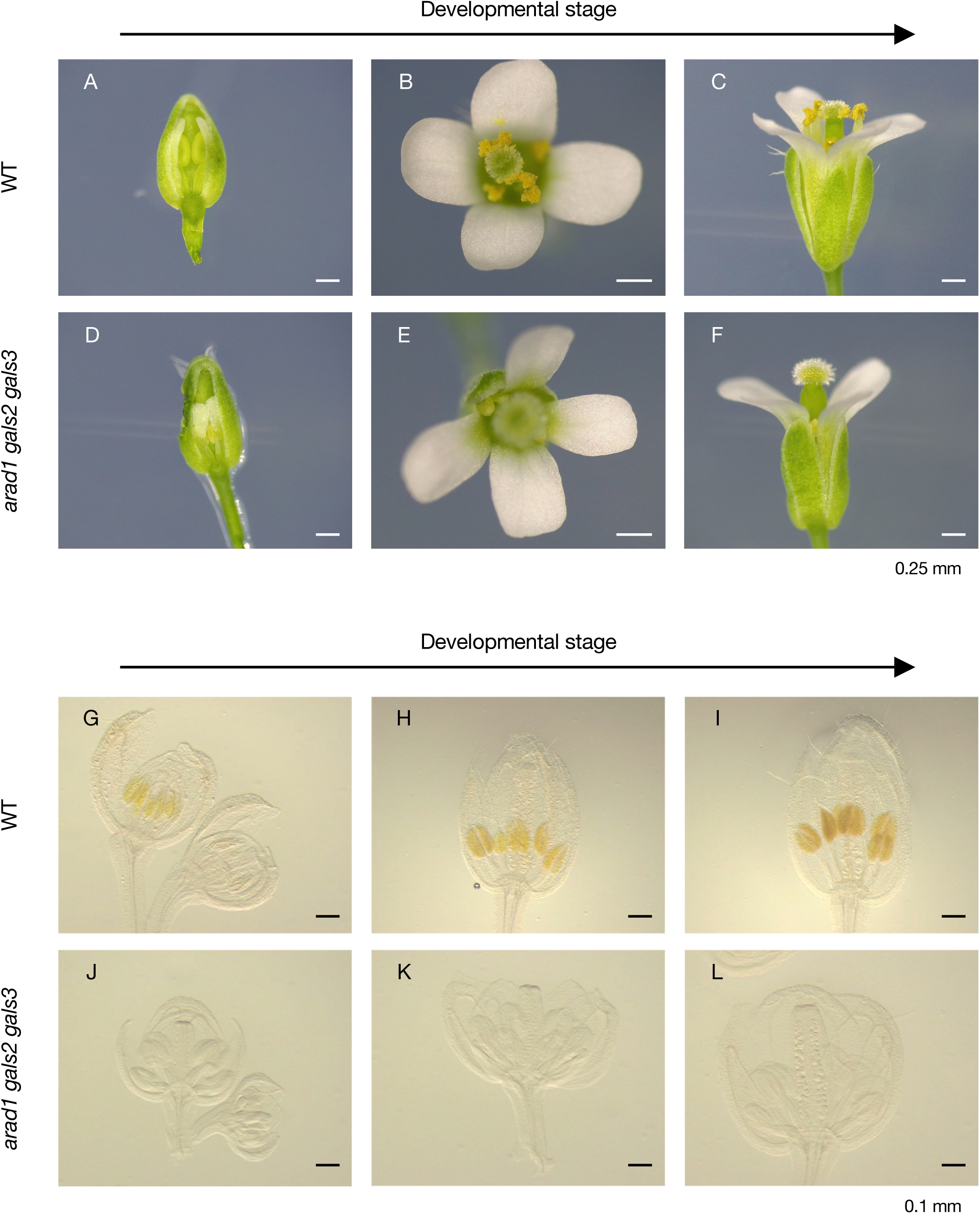
Pollen formation process during the developmental stages of the flower bud observed by optical microscope. (A-C) Development of flower bud morphology in WT. Scale bar = 0.25 mm. (D-F) Development of flower bud morphology in *arad1 gals2 gals3*. Scale bar = 0.25 mm. (G-I) Pollen formation process during the development of flower buds in WT before flowering. Scale bar = 0.1 mm. (J-L) Pollen formation process during the development of flower buds in *arad1 gals2 gals3* before flowering. Scale bar = 0.1 mm. Alt text: (A-L) are images taken with an optical microscope. (A-F) show observations of flower buds in WT and *arad1 gals2 gals3*, either from a cross-section or a top view. These photographs showed that *arad1 gals2 gals3* has short filaments and no pollen formation. (G-L) showed photographs of WT and *arad1 gals2 gals3* buds treated with chloral hydrate for transparency, revealing the process of internal pollen development. These photographs showed the pollen was not clearly formed in the mutant, even before flowering.

To find out if *arad1 gals2 gals3* pollen formation was normal, F1 seeds resulting from pollination of WT pistils with pollen from *arad1/+ gals1/+ gals2 gals3* were sown on medium, and their genotypes were analyzed (Table 1). The pollen genotypes of *arad1/+ gals1/+ gals2 gals3* at the post-meiotic stage were theoretically either *ARAD1 gals1 gals2 gals3* or *arad1 GALS1 gals2 gals3*. The genotypes of the F1 seeds obtained in this experiment showed a nearly one-to-one ratio of *arad1*/+ *GALS1 gals2 gals3* and *ARAD1 gals1*/*+ gals2 gals3* (Table 1). A chi-squared test confirmed that the observed genotype segregation did not deviate from the expected ratio (Table 1). This indicates that pollen with *arad1 gals2 gals3* genotype after meiosis has the ability to fertilize normally. Therefore, we infer these mutations in *ARAD1*, *GALS2* and *GALS3* genes do not affect pollen function after meiosis but instead arrest pollen development during early stages of pollen formation prior to meiosis, resulting in male sterility.

**Table 1.** The relationship between the genotype of pollen after meiosis and male sterility. Pollen collected from plants with *arad1/+ gals1/+ gals2 gals3* genotype was used to pollinate the stigma of WT plants. The genotypes of the resulting F1 population were determined by PCR to identify the pollen genotype from which each seed originated. The differences between the expected and observed segregation ratios of F1 population were statistically analyzed using a chi-squared test. Alt text: Pollen from *arad1/+ gals1/+ gals2 gals3* plants was used to pollinate WT, and the genotypes of the F1 population were analyzed. In *arad1/+ gals1/+ gals2 gals3*, pollen genotypes become either *arad1 GALS1 gals2 gals3* or *ARAD1 gals1 gals2 gals3* after meiosis. Therefore, if pollen with *arad1 gals2 gals3* genotype exhibits abnormalities after meiosis, F1 population derived from *arad1 gals2 gals3* pollen would decrease compared to those from *gals1 gals2 gals3*. Genotyping of F1 population and statistical analysis using a chi-squared test revealed no significant differences, suggesting that *arad1 gals2 gals3* pollen develops normally after meiosis.

| After meiosis pollen genotype | *arad1 gals2 gals3* | *gals1 gals2 gals3* |
| --- | --- | --- |
| Observed | 33 | 38 |
| Expected | 0.5 | 0.5 |
| X-squared | 0.35211 |  |
| *p* -value | 0.5529 |  |

### Arabinan and galactan play an important role in the developmental process of PMCs

To identify developmental defects during the early stage of pollen formation in the mutant, we cut sections from resin-embedded buds and stained the cell walls with toluidine blue (Fig. 3). In WT, pollen development progressed synchronously with the developmental stages of anthers, and pollen tetrads and microspores were observed at stage 7 to stage 8 (Fig. 3A-D). Similarly normal PMCs and tapetum formation in anthers were observed in *arad1 gals2 gals3* as well as WT up to stage 5 (Fig. 3E). However, in *arad1 gals2 gals3*, no tapetums were observed in anthers, and no pollen development to stage 6 or beyond was observed in any individuals examined (Figs. 3F-3H). Instead, a different anther internal structure was observed in *arad1 gals2 gals3* than in the WT (Fig. 3F-D), and the PMCs-like structures appeared to have eventually shrunk (Fig. 3H). This suggests that the accumulation and function of pectic arabinan and galactan around stage 5 is necessary for progression to subsequent stages, such as during PMCs formation and/or the pollen tetrad stage when meiosis is taking place. The findings shown in Table 1 are in agreement with the idea that the genotypes of *ARAD1*, *GALS2* and *GALS3* genes before but not after meiosis affect later seed formation.

**Fig. 3.**
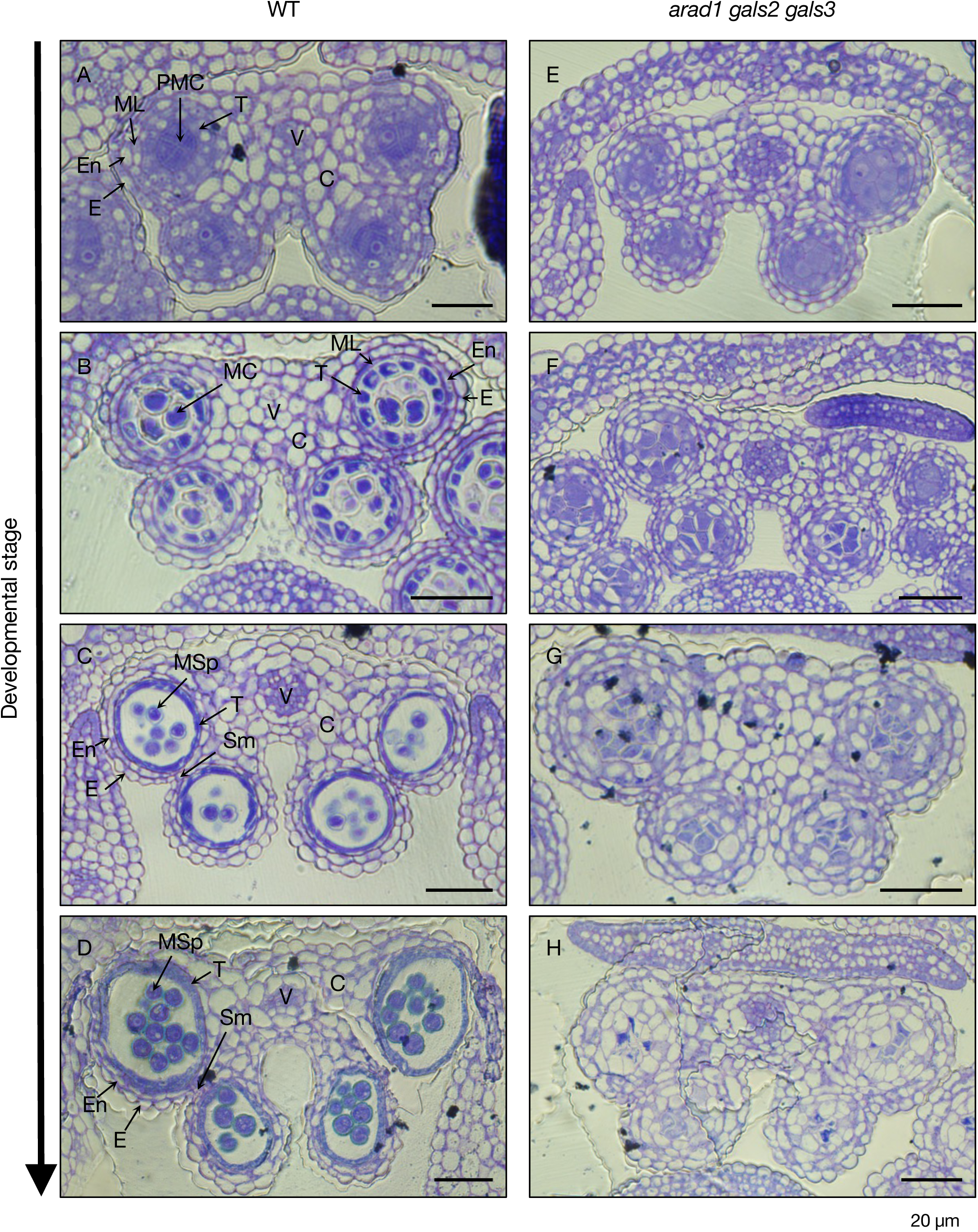
Observation of pollen formation process in WT and *arad1 gals2 gals3* using toluidine blue staining. (A-D) The developmental process of pollen in the WT was observed with toluidine blue staining under an optical microscope, focusing on the progression from meiotic PMCs to microspores. Scale bar = 20 µm. (E-H) The developmental process of pollen in *arad1 gals2 gals3* was observed at different anther developmental stages. Scale bar = 20 µm. C, connective; E, epidermis; En, endothecium; MC, meiotic cell; ML, middle layer; MSp, microspore; PMCs, pollen mother cell; SM, septum; T, tapetum; V, vascular region. Alt text: Images showing the process from PMCs to microspores observed using an optical microscope. These images depict buds embedded and sectioned transversely, stained with toluidine blue to indicate cell wall accumulation sites. In WT, a normal pollen developmental process was observed. By contrast, in *arad1 gals2 gals3*, while PMCs formed at a specific stage (E), subsequent meiotic division did not occur, resulting in abnormal pollen development.

In order to determine the localization of pectin galactan and arabinan accumulation during pollen and anther development, immunohistochemistry was performed (Figs. 4, 5 and Supplementary Fig. S5). Using LM6, an arabinan-specific antibody, arabinan localization was observed during pollen development in WT (Fig. 4), and it was found to accumulate in the cell walls of anther tissues (Fig. 4A-D). Furthermore, arabinan accumulation also occurred at the meristematic surface of the PMCs (Figs. 4E-J), but arabinan disappeared as pollen development progressed (Fig. 4K, L). In tapetum tissues, arabinan disappeared slightly earlier than on the PMCs surface (Fig. 4I, J). Immunohistochemistry with LM5, an antibody specific to pectic galactan, showed a pattern similar to LM6 (Fig. 5). Like arabinan, galactan initially accumulated throughout the anther cells (Fig 5A-F), and then as the PMCs developed, galactan was observed in the meristematic surface of the PMCs while disappearing from the tapetum tissue at that time (Fig. 5G-L). Also, like arabinan, galactan disappeared during the pollen tetrad stage, although it continued to accumulate in other anther cells as pollen developed (Fig. 5C-D, M, N). By contrast, neither arabinan nor galactan was detected in PMCs in *arad1 gals2 gals3* (Fig. 6), but instead a thick cell wall structure, strongly stained with calcofluor white (Fig. 6E, L). This cell wall structure could also be stained with aniline blue, indicating callose accumulation (Fig. 7). The accumulation of callose in PMCs was particularly pronounced during meiosis, and this pattern was observed similarly in both WT and *arad1 gals2 gals3* (Fig. 7A-F). In *arad1 gals2 gals3*, callose continued to accumulate even after the PMCs had failed to develop normally and had collapsed (Fig. 7G, H).

**Fig. 4.**
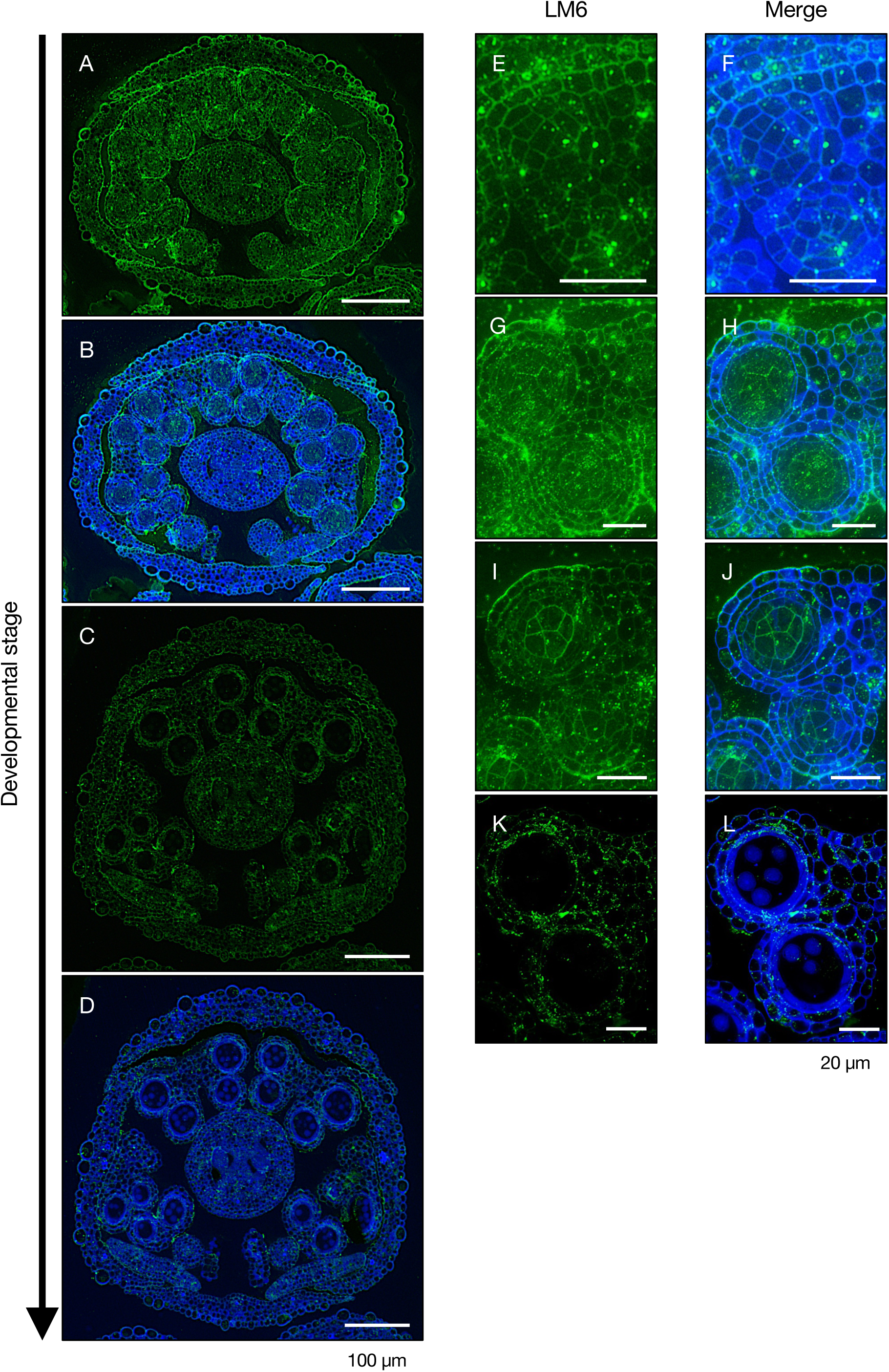
Accumulation of arabinan during pollen development in WT. (A-D) Observation of arabinan distribution in cross sections of an entire flower bud during anther and pollen development. Scale bar = 100 µm. Green-stained regions in (A, C) indicate arabinan accumulation detected with LM6 and Alexa fluor 488 antibodies, while (B, C) show both arabinan accumulation and cell wall visualized with LM6 and calcofluor white. (E-L) Magnified images of the anther showing arabinan accumulation during the pollen development process. Scale bar = 20 µm. (E, G, I, K) Green-stained regions represent Alexa fluor 488 fluorescence of arabinan recognized by LM6. (F, H, J, L) Merged images showed both arabinan accumulation and cell wall regions with LM6 and calcofluor white. Alt text: (A-L) Transverse sections of WT buds embedded and stained, observed using epifluorescence microscope with immunohistochemical staining. (A, C, E, G, I, K) Green staining indicates arabinan accumulation, detected with the LM6 antibody, which specifically recognizes arabinan. (B, D, F, H, J, L) Merged images showing arabinan accumulation with LM6 and cell wall location stained with calcofluor white. In (E, F), no PMCs were observed in the anther, and arabinan accumulation was absent. In (G-J), PMCs began to form in the anther, and arabinan accumulation was detected on the surface of the PMCs, while the PMCs were not stained with calcofluor white. (K, L) Observations of microspores showed no arabinan accumulation, suggesting that arabinan deposition is specific to certain stages of pollen development.

**Fig. 5.**
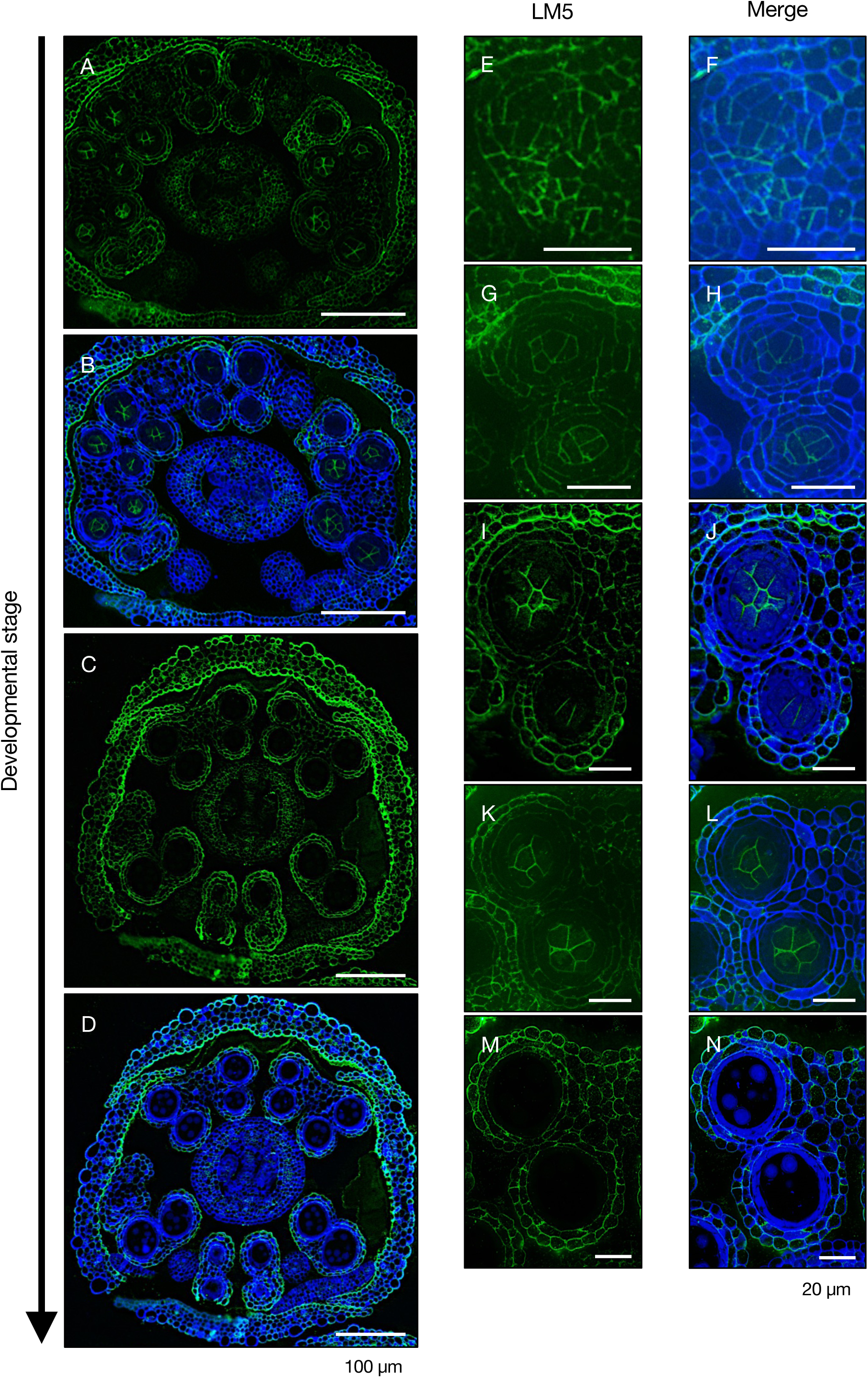
Accumulation of galactan during pollen development in WT. (A-D) Observation of galactan distribution in cross sections of an entire flower bud during anther and pollen development. Scale bar = 100 µm. Green-stained regions in (A, C) indicate galactan accumulation detected with LM5 and Alexa fluor 488 antibodies, while (B, C) show both galactan accumulation and cell wall visualized with LM5 and calcofluor white. (E-L) Magnified images of the anther showing galactan accumulation during the pollen development process. Scale bar = 20 µm. (E, G, I, K, M) Green-stained regions represent Alexa fluor 488 fluorescence of galactan recognized by LM5. (F, H, J, L, N) Merged images showed both galactan accumulation and cell wall regions with LM5 and calcofluor white. Alt text: (A-N) Transverse sections of WT buds embedded and stained, observed using epifluorescence microscope with immunohistochemical staining. (A, C, E, G, I, K, M) Green staining indicates galactan accumulation, detected with the LM5 antibody, which specifically recognizes galactan. (B, D, F, H, J, L, N) Merged images showing galactan accumulation with LM5 and cell wall location stained with calcofluor white. In (E, F), no PMCs were observed in the anther, and galactan accumulation was absent. In (G-J), PMCs began to form in the anther, and galactan accumulation was detected on the surface of the PMCs, while the PMCs itself were not stained with calcofluor white. (K, L, M, N) Observations of microspores showed no galactan accumulation, suggesting that galactan deposition is specific to certain stages of pollen development.

**Fig. 6.**
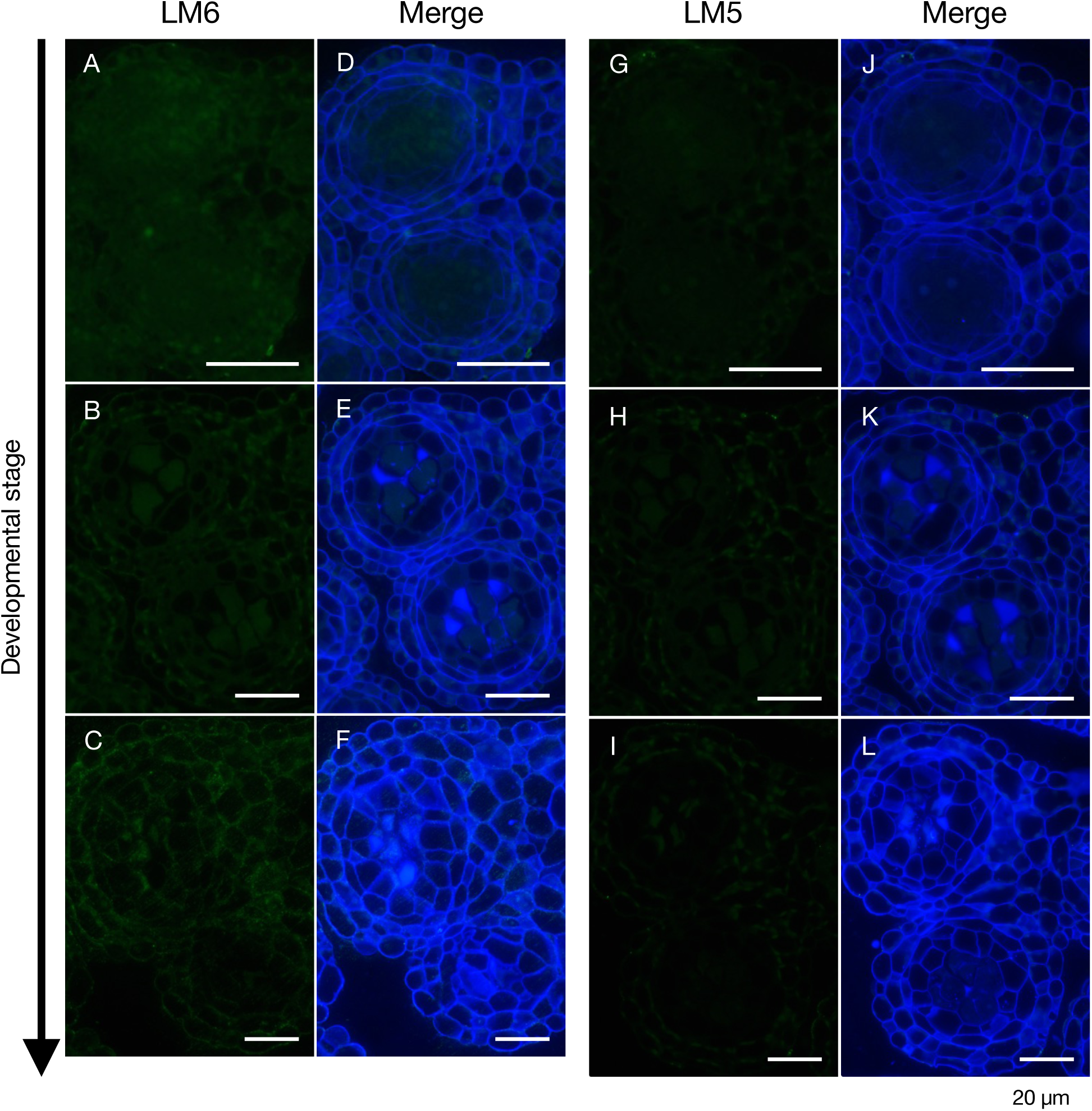
Accumulation of arabinan and galactan during pollen development in *arad1 gals2 gals3*. The accumulation of arabinan (A-F) and galactan (G-L) during the pollen development process in the anther of *arad1 gals2 gals3* was observed by immunohistochemical staining. Scale bar = 20 µm. Arabinan (A-C) and galactan (G-I) accumulation was detected using the LM6 and LM5 antibody, respectively. (D-F, J-L) Merged images of calcofluor white and LM6 (D-F) and LM5 (J-L). Alt text: Transverse sections of *arad1 gals2 gals3* buds embedded and stained, observed using epifluorescence microscope with immunohistochemical staining. (A-F) Arabinan accumulation was visualized using antibody LM6, which specifically recognized arabinan, but in contrast WT, no accumulation was observed in *arad1 gals2 gals3*. (G-L) The same was observed with galactan staining using LM5. (E, L) When the cell wall was visualized using calcofluor white, some cell wall polysaccharides accumulated in the PMCs were observed in *arad1 gals2 gals3*. (F, L) As development progressed, the PMCs were observed to lose its shape.

**Fig. 7.**
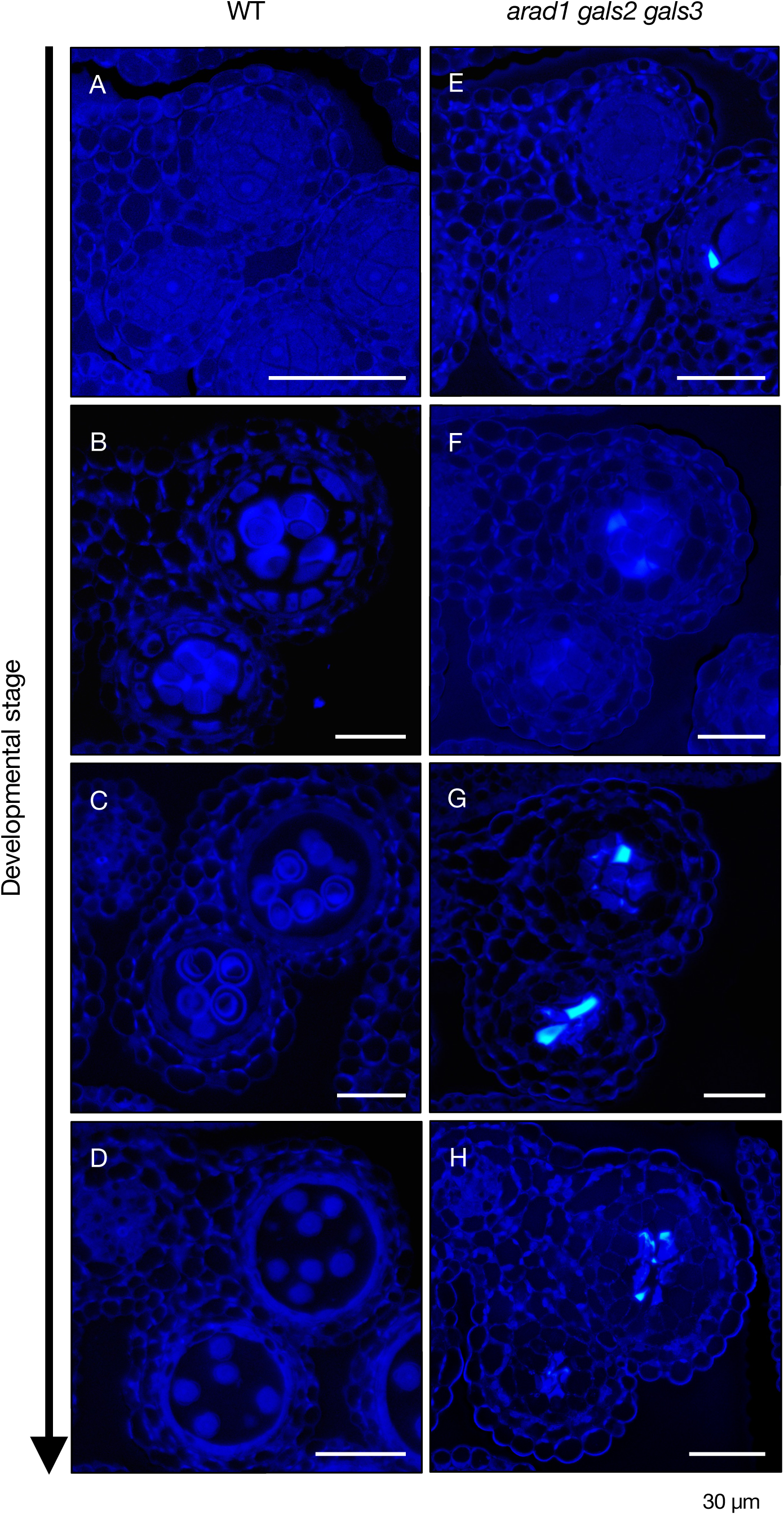
The accumulation of callose during the pollen development process in WT and *arad1 gals2 gals3*. (A-H) The accumulation of callose during the pollen development progress was observed using aniline blue staining in WT (A-D) and *arad1 gals2 gals3* (E-H). Scale bar = 30 µm. Alt text: Transverse sections of WT and *arad1 gals2 gals3* buds embedded and observed using epifluorescence microscope with aniline blue staining to examine callose accumulation. (A, E) Callose accumulation in PMCs showed minimal differences between WT and *arad1 gals2 gals3* during early pollen development. (B-D) In WT, callose accumulated on the surface of pollen following meiosis. (F) In *arad1 gals2 gals3*, callose began to accumulate before the meiotic stage. (G, H) After meiosis, callose was found to accumulate on the surface of significantly distorted pollen. These observations revealed that even in *arad1 gals2 gals3*, where pollen failed to develop normally, hcallose still accumulated on the surface of the pollen.

## Discussion

This study demonstrates that arabinan and galactan are crucial for normal pollen development. Previous research has indicated that significant reductions in arabinan in *arad1* or galactan in *gals1 gals2 gals3* did not notably affect pollen development or plant growth (Harholt *et al*., 2006, 2012; Liwanag *et al*., 2013). However, in this study, the simultaneous reduction of arabinan and galactan in *arad1 gals2 gals3* showed minimal effects on growth but resulted in male sterility, preventing seed production. A look at mRNA expression retrieved from the eFP browser (https://bar.utoronto.ca/efp/cgi-bin/efpWeb.cgi) revealed that *ARAD1*, *GALS2* and *GALS3* are expressed in various tissues, including stamens (Supplementary Fig. S6-8). Interestingly, *arad1 gals2* and *arad1 gals3* double mutants also partially failed to produce seeds, suggesting that the presence of sufficient arabinan and galactan, rather than the specific genes, is critical for normal pollen formation and seed production (Supplementary Fig. S9). By contrast, other male sterility mutants like *ap1*, *ap2*, and *lug* exhibit additional phenotypes such as floral organ abnormalities or altered leaf morphology (Irish and Sussex, 1990; Jofuku *et al*., 1994; Shi *et al*., 2024). This distinction points to a unique mechanism underlying the seed production failure in *arad1 gals2 gals3*. These findings underline the importance of arabinan and galactan in seed production and suggest that ARAD1, GALS2 and GALS3 play a vital role in synthesizing these polysaccharides, especially during pollen development.

The inability of *arad1 gals2 gals3* to produce seeds appears to be due to the failure of pollen development (Fig. 2, 3). It is known that cell wall components are involved in pollen maturation (Dong *et al*., 2005; Enns *et al*., 2005; Cankar *et al*., 2014; Hasegawa *et al*., 2023), particularly pectins, where the methylesterification of HG in *Oryza sativa* and arabinan in potato has been shown to be essential for normal pollen formation (Cankar *et al*., 2014; Hasegawa *et al*., 2023). However, in Arabidopsis, *arad1* with reduced arabinan and *gals1 gals2 gals3* with reduced galactan can produce seeds similarly to the WT. Thus, the role of arabinan and galactan in pollen development was largely unknown. This study demonstrates that both arabinan and galactan are important for the early development of pollen prior to meiosis in Arabidopsis and it is crucial that at least one is available. While previous research has shown that callose is also critical for normal pollen development (Dong *et al*., 2005; Enns *et al*., 2005; Chen and Kim, 2009), a certain level of callose accumulation occurred even after a stage in which no accumulation of arabinan and galactan was observed in the *arad1 gals2 gals3* mutant (Fig. 7). These results suggest that arabinan and galactan play key roles during PMCs meiosis, and their joint reduction causes male sterility independently of callose accumulation.

Arabinan and galactan are neutral sugar side chains of pectin and it has been suggested that they bind to cellulose microfibrils (Lin *et al*., 2015). This interaction is thought to enhance cell wall strength and create appropriate spacing between pectin main chains including HGs (Moore *et al*., 2008; Verhertbruggen *et al*., 2009; Harholt *et al*., 2010; Mariette *et al*., 2021). In this study, arabinan and galactan were observed to accumulate at the cell adhesion surfaces of PMCs. These components are deposited in the primary cell wall, but they may also accumulate in the middle lamella. The middle lamella, primarily composed of pectin, acts as an adhesive layer between adjacent primary cell walls (Zamil and Geitmann, 2017). During the synthesis of HG in the Golgi apparatus, it is methyl esterified (Wolf *et al*., 2009; Wolf and Greiner, 2012), but its demethyl-esterification, facilitated by pectin methylesterase (PME), plays a critical role in the adhesive properties of the pectin and thus of the middle lamella (Bosch and Hepler, 2005; Pelloux *et al*., 2007). Demethyl-esterified HGs, with nine or more consecutive de-methylated GalA units, can form calcium-mediated cross-links, leading to gelation (Wolf *et al*., 2009). In Arabidopsis, PME has also been implicated in the separation of PMCs into a tetrad post-meiosis (Francis *et al*., 2006). This separation process requires demethyl-esterified HGs, which can be cleaved by endo-polygalacturonases (PGs) (Wolf *et al*., 2009). HG cleavage is effective only when it has undergone adequate demethyl-esterification (Wolf *et al*., 2009; Sénéchal *et al*., 2014). Taking findings of previous studies and of this one into consideration, the abnormal meiotic process observed in *arad1 gals2 gals3* may result from the reduced levels of arabinan and galactan, leading to the closer proximity of HG (Moore *et al*., 2008; Harholt *et al*., 2010; Mariette *et al*., 2021). This proximity may disrupt proper dimethyl-esterification, which is critical for normal meiotic progression.

Furthermore, it has been suggested that the reduction of arabinan and galactan may affect not only pollen development but also other organs of the floral bud. This study found that, in *arad1 gals2 gals3*, filaments were shorter and potentially underdeveloped compared to the WT (Fig. 2). This suggests that either the development of pollen or anthers affects filament growth or that the accumulation of arabinan and galactan may play an essential role in filament development. To date, there have been no reports linking pollen or anther development with filament growth, nor studies suggesting a relationship between arabinan and galactan accumulation and filament development. Therefore, we hypothesize that the development of filaments and anthers may be synchronized. These findings could provide new insights into the unexplored aspects of reproductive tissue development in plants.

In conclusion, this study revealed that arabinan and galactan work together, particularly in pollen meiosis while having no apparent effect on ovule development, at least in Arabidopsis. While it remains to be determined whether arabinan and galactan are universally important for all plants undergoing sexual reproduction, these polysaccharides are consistently found in the cell walls of organisms ranging from bryophytes to seed plants (Harholt *et al*., 2010; Silva *et al*., 2011; Roberts *et al*., 2012). Moreover, the *ARAD* and *GALS* genes, which are responsible for synthesizing arabinan and galactan, are also conserved in land plants. They are major components of the cell walls in the aerial parts of angiosperms, particularly in dicotyledons (Lin *et al*., 2015). The involvement of pectin in pollen development has been widely reported in various angiosperms including Arabidopsis, which was used in this study (Cankar *et al*., 2014; Hasegawa *et al*., 2023). By contrast, the composition of pollen cell walls in gymnosperms is less consistent than in angiosperms, with the pectin content in pollen varying by species, and some species reportedly lacking pectin altogether (Yatomi *et al*., 2002; Harholt *et al*., 2010; Breygina *et al*., 2021). Based on these findings, it is likely that the importance of pectin, including arabinan and galactan, in pollen development was conserved during the evolutionary process following the emergence of angiosperms.

## Abbreviations

ARAD: ARABINAN DEFICIENT
C: connective
E: epidermis
En: endothecium
GALS: GALACTAN SYNTHASE
HG: homogalacturonan
MC: meiotic cell
ML: middle layer
MSp: microspores
PMCs: pollen mother cells
RG: rhamnogalacturonan
SM: septum
T: tapetum
V: vascular region

## Supplementary data

Table S1. Primer sequences used in this study.

Table S2. Past research on the relationship between pollen development and cell wall components.

Fig. S1. Pectin content, monosaccharide composition ratio, and monosaccharide content in the aboveground parts of WT and *arad1 gals2 gals3*.

Fig. S2. Cell wall extensibility and breaking force of the rosette leaves of WT and *arad1 gals2 gals3*.

Fig. S3. Histogram of the number of seeds per silique in *gals1 gals2 gals3*, *arad1/+ gals1/+ gals2 gals3*, and *arad1 gals2 gals3*.

Fig. S4. The genotype of F1 seeds obtained by crossing WT pollen with *arad1 gals2 gals3* mutant.

Fig. S5. Control groups for immunohistochemical observations of pollen development in WT and *arad1 gals2 gals3* mutants.

Fig. S6. Expression atlas of the Arabidopsis *ARAD1* gene across various tissues, retrieved from the eFP Browser.

Fig. S7. Expression atlas of the Arabidopsis *GALS2* gene across various tissues, retrieved from the eFP Browser.

Fig. S8. Expression atlas of the Arabidopsis *GALS3* gene across various tissues, retrieved from the eFP Browser.

Fig. S9. Phenotype of *arad1 gals2* and *arad1 gals3*.

## Acknowledgement

We are grateful to Profs. Kenichi Nonomura (National Institute of Genetics) and Sumie Ishiguro (Nagoya University) for their critical assistance.

## Author contributions

Conceptualization, T, Kikuchi, T. Kotake and D. T.; investigation, T. Kikuchi, K. S., T. Kotake and D. T.; writing – original draft, T, Kikuchi and D. T.; writing – review & editing, T. Kotake and D. T.; supervision, T. Kotake and D. T.; funding acquisition, T. Kotake and D. T.

## Data availability

Data are available from the corresponding author upon request.

